# Transcriptomic interplay between *Acinetobacter baumannii*, human macrophage and polymyxin

**DOI:** 10.1101/2024.01.23.576770

**Authors:** Zhi Ying Kho, Mohammad Abul Kalam Azad, Yan Zhu, Mei-Ling Han, Qi (Tony) Zhou, Tony Velkov, Thomas Naderer, Jian Li

## Abstract

Optimization of antibiotic therapy has been hindered by our dearth of understanding on the mechanism of the host-pathogen-drug interactions. Here, we employed dual RNA-sequencing to examine transcriptomic perturbations in response to polymyxin B in a co-culture infection model of *Acinetobacter baumannii* and human macrophages. Our findings revealed that polymyxin B treatment induced significant transcriptomic response in macrophage-interacting *A. baumannii*, exacerbating bacterial oxidative stress, disrupting metal homeostasis, affecting osmoadaptation, triggering stringent stress response, and influencing pathogenic factors. Moreover, infected macrophages adapt heme catabolism, coagulation cascade, and hypoxia-inducible signaling to confront bacterial invasion. Disrupting *rcnB*, *ompW*, and *traR/dksA* genes in *A. baumannii* impairs metal homeostasis, osmotic stress defense and stringent responses, thereby enhancing antibacterial killing by polymyxin. These findings shed light on the global stress adaptations at the network level during host-pathogen-drug interactions, revealing promising therapeutic targets for further investigation.

**IMPORTANCE:** In the context of the development of bacterial resistance during the course of antibiotic therapy, the role of macrophages in shaping bacterial response to antibiotic killing remains enigmatic. Herein we employed dual RNA-sequencing and an *in vitro* tripartite model to delve into the unexplored transcriptional networks of the *Acinetobacter baumannii*-macrophage-polymyxin axis. Our findings uncovered the potential synergy between macrophages and polymyxin B which appear to act in co-operation to disrupt multiple stress tolerance mechanisms in *A. baumannii*. Notably, we discovered the critical roles of bacterial nickel/cobalt homeostasis (*rcnB* family), osmotic stress defense (*ompW* family), and stringent response regulator (*traR/dksA* C4-type zinc finger) in tolerating the last-line antibiotic polymyxin B. Our findings may lead to potential targets for the development of novel therapeutics against the problematic pathogen *A. baumannii*.

## INTRODUCTION

In recent decades, the evolution of life-threatening infections caused by Gram-negative bacterial ‘superbugs’ has exceeded the pace at which we are developing novel classes of antibiotic therapeutics [1, 2]. Multi-drug resistant (MDR) *Acinetobacter baumannii* is a particularly difficult and is a leading cause of nosocomial infections worldwide [3]. Designated by the World Health Organization (WHO) as a “Priority 1: Critical” pathogen, carbapenem-resistant *A. baumannii* urgently necessitates novel therapeutic options [4–6]. Polymyxins (i.e., polymyxin B and colistin) are lipopeptide antibiotics that are employed as a last-line therapy for many MDR Gram-negative infections, such as those caused by *A. baumannii* [7]. Worryingly, the suboptimal use of polymyxins has facilitated the emergence of polymyxin resistance, including in *A. baumannii* [8, 9]. With its high genomic and metabolic plasticity, *A. baumannii* can adapt to diverse environmental stressors, including the host immune system and antibiotics [10–17]. Therefore, an in-depth comprehension of the intertwined molecular adaptations associated with infection and drug tolerance is imperative for discovering novel drug targets and developing innovative immunological therapeutics.

Current antibiotic pharmacodynamic research usually overlooks the impact of the host environment on bacterial adaptability [18]. Herein we elucidated the interspecies transcriptional responses during different treatment conditions using a tripartite model consisting of MDR *A. baumannii* AB5075 (a model MDR strain), THP-1 human macrophage, and polymyxin antibiotic. Using a transposon mutant library, we revealed the indispensable role of metal homeostasis, osmoadaptation and stringent response in bacterial tolerance to polymyxins. Furthermore, hypoxia-inducible signaling and iron-heme homeostasis in THP-1 human macrophages have emerged as pivotal players in countering *A. baumannii* invasion.

## RESULTS

### *A. baumannii* transcriptional responses to macrophage, polymyxin B and in combination

Herein we employed dual RNA-sequencing to investigate global responses in the macrophage-*A. baumannii*-polymyxin B axis (**Fig S1**). We uncovered 3,744 bacterial transcripts across the bacteria-associated samples. The interacting bacteria (adherent and intracellular) from the THP-1 human macrophage (THP) infection group (*A. baumannii* [AB]+THP) displayed the most distinct separation from the untreated *A. baumannii* controls in the first component of the principal component analysis (PCA) plot, followed by the polymyxin B (PMB) treated infection group (AB+THP+PMB); and the bacteria treated solely with polymyxin B (AB+PMB) (**Fig S2A**). *A. baumannii* in the presence of macrophage exhibited 1,126 statistically significant differentially expressed genes (DEGs), compared to *A. baumannii* alone (log_2_ fold change [FC] >1 or <-1, and false discovery rate [FDR] adjusted p-value <0.05), followed by AB+THP+PMB *vs*. AB (764 DEGs) and AB+PMB *vs*. AB (435 DEGs). These findings underscore the profound transcriptional perturbations in *A. baumannii* arising from bacterial interaction with macrophage (**Fig S2B, Table S1-S3**).

#### Macrophage reshapes A. baumannii redox, osmotic stress response, and virulence landscape

The macrophage infecting *A. baumannii* AB5075 cells overexpressed several redox stress tolerance genes, including cold shock protein *csp1* (ABUW_RS13055; log_2_FC, 1.66), glutaredoxin *grxC* (log_2_FC, 1.58), glutathione *S*-transferase (ABUW_RS19900; log_2_FC, 1.01), alkyl hydroperoxide reductase *ahpF* (log_2_FC, 1.30) and RNA polymerase-binding protein encoded by *dksA* (log_2_FC, 1.00); compared to the untreated bacterial controls (**Fig 1A**). Moreover, interacting bacteria from the AB+THP group upregulated multiple ABC transport and assimilation systems responsible for sulfate, involving the cysteine regulon *cysTPWD* and *sbp* (ABUW_RS01375), as well as taurine, a sulfur-containing amino acid, facilitated by *tauACD* and FMNH2-dependent alkanesulfonate monooxygenase *ssuD* (**Fig 1B**). Conversely, several thiol-based antioxidant genes were downregulated in *A. baumannii* AB5075 in the presence of macrophages, compared to the untreated control group, encompassing multiple glutathione *S*-transferases, lactoylglutathione lyase *gloA*, glutathione peroxidases (*gpo* and *btuE*), thioredoxin *trxC* and thioredoxin-disulfide reductase *trxB* (**Fig 1A**).

**Figure 1:**
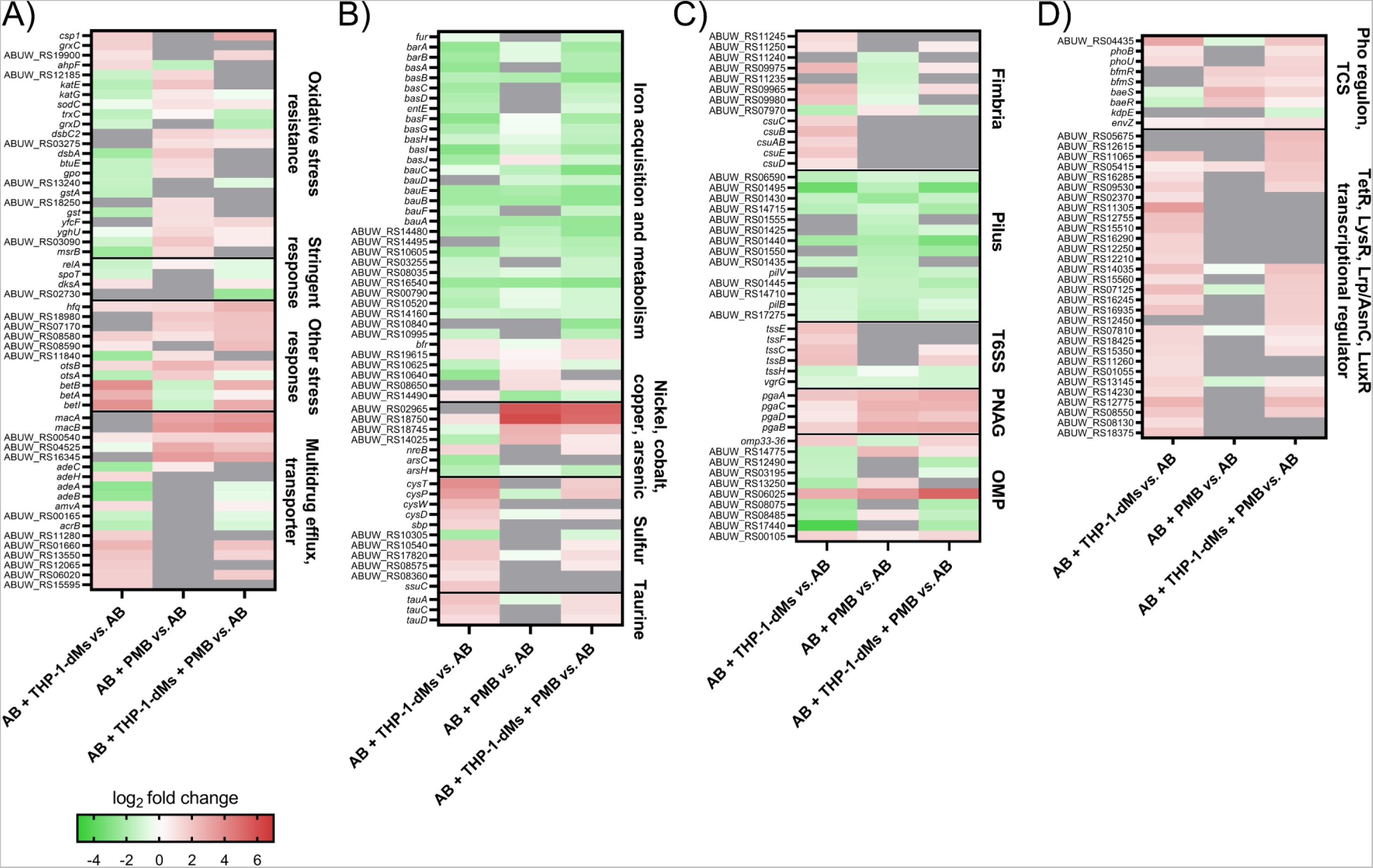
Transcriptomic signatures of *A. baumannii* AB5075 under different selective pressures. Heatmaps delineating expression of bacterial genes from the categories of **A**) stress response and multidrug efflux; **B**) homeostases of metals and other metabolites; **C**) pathogenic factors; **D**) signal transduction and transcriptional regulators in respective comparison groups. The colour code shows log_2_FC values of gene expression with FDR <0.05 compared to that of untreated bacteria. Grey represents gene expression with FDR ≥0.05. Data are presented as the mean of biological triplicates per group. AB, *A. baumannii* AB5075; THP, THP-1 differentiated macrophages; PMB, polymyxin B; TCS, two-component system.

Notably, interacting bacteria from the AB+THP group exhibited significant overexpression of osmoadaptation genes, including trehalose phosphatase *otsB* (log_2_FC, 1.29), choline dehydrogenase *betA* (log_2_FC, 2.62), betaine-aldehyde dehydrogenase *betB* (log_2_FC, 3.62), and *ompW* family ABUW_RS06025 (log_2_FC, 2.81) (**Figs 1A, 1C**). Interacting bacteria from the AB+THP group also exhibited a distinctive upregulation of virulence-associated fimbrial biogenesis genes and the Type VI secretion system (T6SS) (**Fig 1C**).

#### Polymyxin B treatment perturbs bacterial cell envelope, redox and metal homeostasis

Following polymyxin B treatment, *A. baumannii* in the AB+PMB group underwent dynamic gene expression changes, particularly in membrane-related components. Noteworthy alterations included the overexpression of lipoprotein (Lol) system genes (*lolABCD*), and maintenance of lipid asymmetry (*mlaBD*; **Fig 2**). Furthermore, bacterial cell envelope stress-sensing and signal transduction two-component systems (TCS) *baeSR* and *bfmSR* were overexpressed in response to polymyxin B treatment (**Fig 1D**). Additionally, compared to the untreated controls a significant overexpression of multidrug efflux systems was observed in the AB+PMB group, including *macAB*, *adeC/adeK/oprM* family multidrug efflux complex outer membrane factor (ABUW_RS16345), RND transporter permease subunit ABUW_RS04525 and MATE family efflux transporter ABUW_RS00540 (**Fig 1A**).

**Figure 2:**
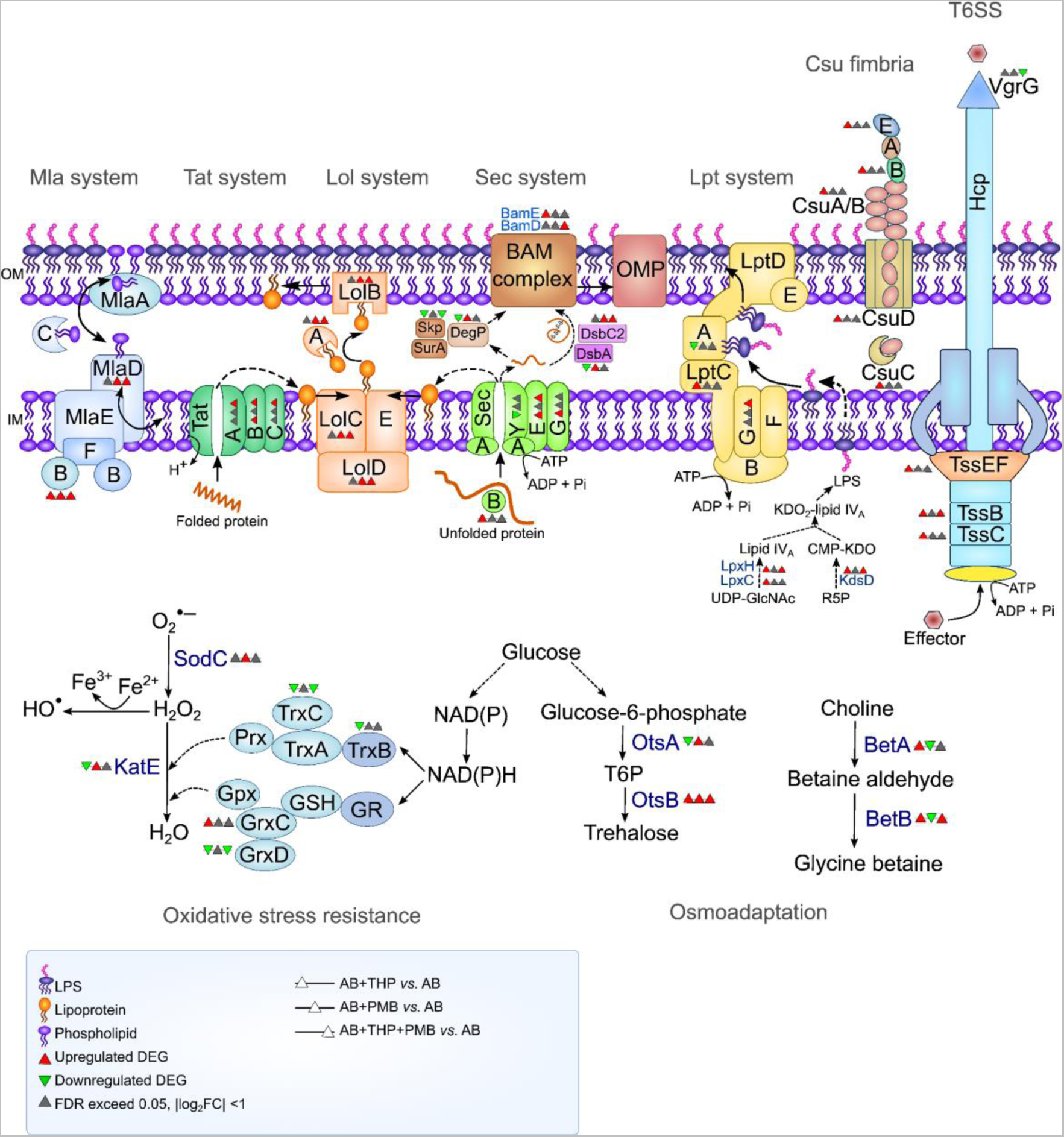
*A. baumannii* AB5075 differentially regulates its secretion and stress adaptation systems in response to macrophages and polymyxin B challenge. Data are presented as the mean of biological triplicates per group. AB, *A. baumannii* AB5075; THP, THP-1 differentiated macrophages; DEGs, differentially expressed genes; GR, glutathione reductase; GSH, glutathione; Gpx, glutathione peroxidase; Prx, peroxiredoxin; T6P, trehalose-6-phosphate.

Another major finding of our study is that polymyxin B treatment alone caused the upregulation of distinct bacterial redox stress resistance systems, separate from those observed in the AB+THP group (**Fig 1A**). This response included the overexpression of reactive oxygen species (ROS) scavengers, such as catalase *katE*, catalase family peroxidase ABUW_RS12185 and superoxide dismutase *sodC*, as well as various thiol-based antioxidants, including glutathione peroxidases, glutathione *S*-transferases, glutathione-dependent disulfide-bond oxidoreductase *yghU* and peroxiredoxin ABUW_RS03090 (**Fig 1A**). Furthermore, polymyxin B treatment induced transcriptional upregulation of bacterial protein folding chaperone systems, encompassing thiol:disulfide interchange protein *dsbA*, *dsbC* family, and peptide-methionine (R)-*S*-oxide reductase *msrB* in the AB+PMB group compared to the untreated control (**Fig 1A**). Intriguingly, polymyxin-treated AB5075 exhibited remarkable overexpression of genes from the nickel/cobalt homeostasis-associated *rcnB* family, including ABUW_RS18750, ABUW_RS02965 and ABUW_RS18745, with respective log_2_FC values of 6.09, 5.74 and 2.63 (**Fig 1B**).

#### Polymyxin B and macrophages induce cooperative biochemical turmoil in the bacterial stress response and pathogenic gene expression

During polymyxin-treated macrophage infection (AB+THP+PMB), interacting bacteria overexpressed several key stress resistance genes. Cold shock protein *csp1*, envelope stress associated RNA chaperone *hfq* and osmotic stress-associated *ompW* family ABUW_RS06025 were notably upregulated (log_2_FC = 2.74, 2.59 and 5.24, respectively; [**Figs 1A, 1C**]). Intriguingly, universal stress protein-coding ABUW_RS08590 and *envZ* (ABUW_RS01255, a global sensor of acidic pH and osmotic stress) exhibited exclusive upregulation in the tripartite condition (AB+THP+PMB), with the respective log_2_FC values of 2.23 and 1.07 compared to the untreated bacteria control (**Figs 1A, 1D**). In the absence of polymyxin B, the interacting bacteria overexpressed redox stress resistance genes (*grxC* and *ahpF*) in the presence of macrophage (AB+THP); however, under the tripartite conditions (AB+THP+PMB) their expressions decreased to the levels approximating the bacteria alone control (**Fig 1A**). Noteworthily, the expression levels of *katE*, catalase family peroxidase ABUW_RS12185, *dsbA*, glutathione peroxidases (*gpo* and *btuE*), *yghU*, glutathione *S*-transferases (ABUW_RS18250 and *gst*) and peroxiredoxin ABUW_RS03090, were upregulated in the AB+PMB group, but returned to the levels similar to those of untreated bacteria in the presence of macrophage under the tripartite condition (**Fig 1A**). Interestingly, stringent stress response-associated ABUW_RS02730 (*traR/dksA* C4-type zinc finger) was exclusively downregulated under tripartite conditions.

Another notable discovery from this study is that exposure to macrophages and polymyxin B, alone or in combination, led to the downregulation of multiple iron acquisition and assimilation genes in *A. baumannii*. The *bas-bau-bar* gene cluster, energy transducer *tonB* (ABUW_RS14480) and TonB-dependent receptors (ABUW_RS10605, ABUW_RS16540, ABUW_RS14160) were generally suppressed (**Fig 1B**). Remarkably, a tripartite-specific downregulation of the *motA/tolQ/exbB* proton channel family (ABUW_RS10840; log_2_FC, −2.62) was observed (**Fig 1B**). Intriguingly, exposure to THP-1 induced overexpression of bacterial virulence factors, such as the *csu* fimbrial operon and *tssCEF* from Type VI secretion system (T6SS), which were subsequently reduced by polymyxin B in the tripartite condition, approaching levels comparable to those of untreated bacteria (**Fig 1C**). ABUW_RS00575, which encodes for the T6SS tip protein VgrG, was exclusively downregulated in the tripartite condition (**Fig 1C**). Moreover, treatment with macrophage and polymyxin B alone (AB+THP or AB+PMB) or in combination (AB+THP+PMB) induced overexpression of the AB5075 *pga* operon which is responsible for poly-β-1,6-*N*-acetyl-D-glucosamine (PNAG) extracellular matrix biosynthesis and crucial for biofilm production (**Fig 1C**). In short, tripartite conditions induced membrane-associated stress in bacteria and cooperative suppression by macrophages and polymyxin B affected bacterial oxidative stress defense, iron uptake, and the virulence-associated T6SS. Treatments with macrophages and/or polymyxin B prompted *A. baumannii* to transcriptionally attempt biofilm formation as a response to environmental stressors.

### Macrophage transcriptional responses to *A. baumannii* AB5075 infection and polymyxin treatment

A total of 21,058 human transcripts were detected across all macrophage samples and macrophage infected with *A. baumannii* AB5075 (AB+THP) exhibited the most distinct separation from untreated macrophage controls in the first component of the PCA plot (**Fig S2D**). Host expression profiles of polymyxin-treated infection (AB+THP+PMB) closely clustered with those of infection alone (AB+THP) (**Fig S2D**). Polymyxin B treatment alone at 30 mg/L without bacterial infection (THP+PMB) closely clustered with untreated host controls (**Fig S2D**). In the AB+THP group, 5,164 statistically significant DEGs were observed compared to the untreated macrophage control. Treatment with 30 mg/L polymyxin B alone for 1 h (THP+PMB) did not yield any detectable DEGs when normalized against untreated macrophage. These findings confirm the absence of observable toxicity of polymyxin (30 mg/L) or its immunomodulatory effect at the host transcriptional level. Consequently, the focus of subsequent analyses centered on the transcriptomic perturbations observed in infected macrophage (AB+THP *vs.* THP).

#### Macrophage orchestrates immune response, coagulation activation and iron-heme modulation during A. baumannii infection

In infected macrophages, a number of pathogen-sensing pattern recognition receptors (PRRs) and their signaling pathways were upregulated, including Toll-like receptor 2 (TLR2; log_2_FC, 1.21), NOD-like receptors, C-type lectin receptors, and RIG-I-like receptors (**Figs 3**, **S2E**). Concomitantly, the expression of genes encoding pro-inflammatory cytokines such as tumor necrosis factor (TNF; log_2_FC, 9.92) and interleukins IL1A, IL1B and IL6 (log_2_FC values of 5.88, 3.95 and 5.05, respectively) was markedly upregulated (**Table S4**). Upregulated DEGs in infected macrophages were significantly enriched in the TNF signaling pathway (**Fig S2E**, **Table S4**). Notably, coagulation cascade genes, such as tissue factor F3 (log_2_FC, 3.26), and tissue plasminogen activator (tPA) inhibitors SERPINB2 (log_2_FC, 3.28) and SERPINE1 (log_2_FC, 2.50), were also upregulated in infected macrophages, highlighting their involvement in the host response (**Fig 3**).

**Figure 3:**
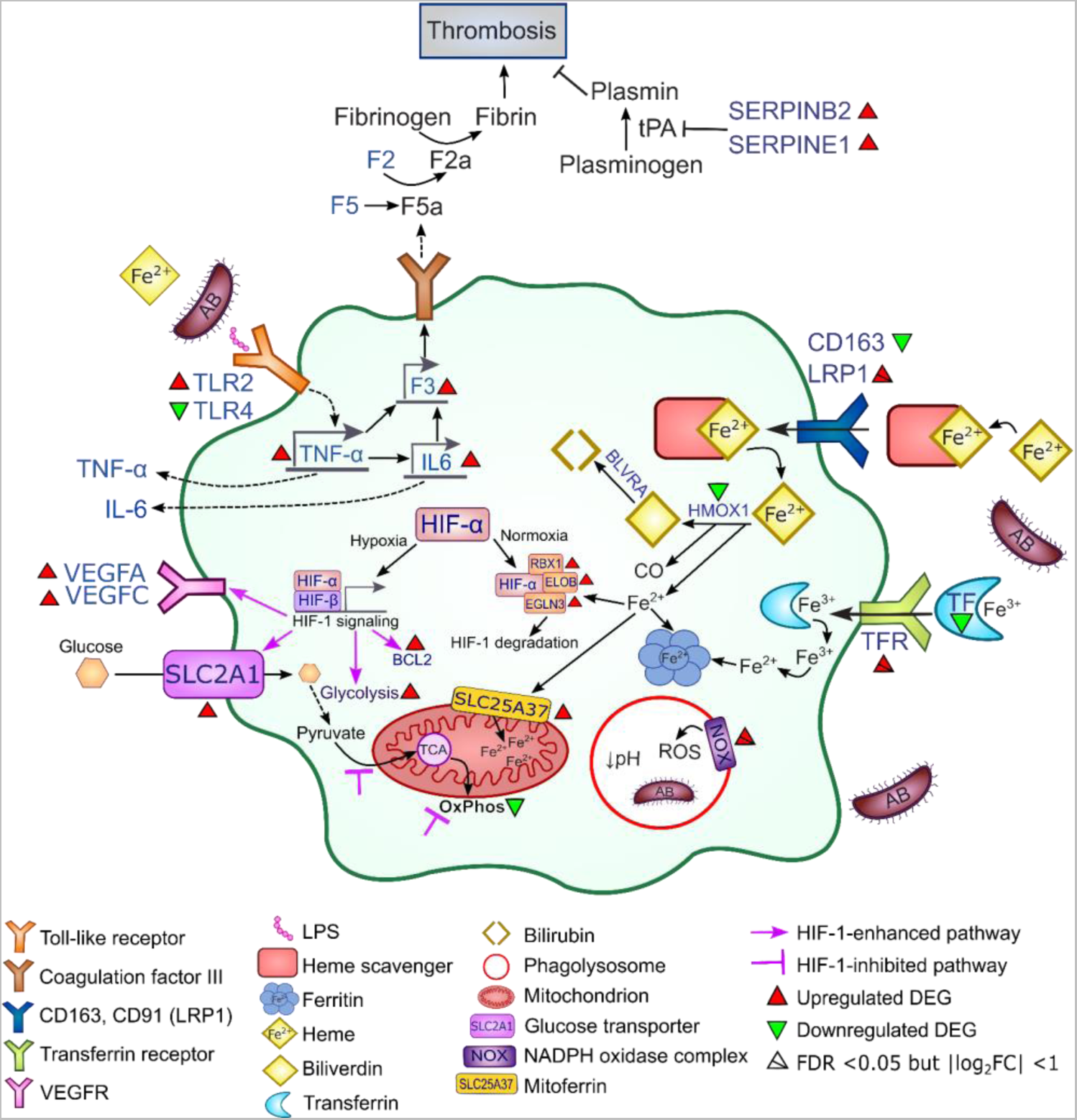
*A. baumannii* AB5075 infection-induced transcriptional modulation of heme catabolism, coagulation cascade and hypoxia-inducible (HIF-1) signaling in macrophages (AB+THP *vs*. THP). Data are presented as the mean of biological triplicates per group. AB, *A. baumannii* AB5075; THP, THP-1 differentiated macrophages; DEGs, differentially expressed genes.

Host iron-heme homeostasis was modulated during infection, as indicated by the upregulation of host metal transporter SLC39A14 (log_2_FC, 2.28). Despite not surpassing the statistical significance cut-off, transferrin receptor protein 1 (TFRC; log_2_FC, 0.996; FDR, 2.24х10^-9^), transferrin receptor protein 2 (TFR2; log_2_FC, 1.674; FDR, 0.092) and ferritin heavy chain (FTH1; log_2_FC, 0.97; FDR, <0.05) exhibited increased expression in infected macrophages (**Fig 3, Table S4**). Moreover, the mitochondrial iron influx-associated host mitoferrin-1 (SLC25A37; log_2_FC, 2.35) was significantly upregulated, while serotransferrin (TF; log_2_FC, −1.67) and heme catabolism associated heme oxygenase I (HMOX1; log_2_FC, −1.80) were both downregulated (**Fig 3**).

#### A. baumannii infected macrophage dynamically rewire hypoxia-inducible signaling

Following infection with *A. baumannii* AB5075, macrophages exhibited pronounced upregulation of genes associated with hypoxia-inducible transcription factor-1 (HIF-1) signaling (**Figs 3**, **S2E**). This was evident through the increased expression of vascular endothelial growth factors (VEGFA, VEGFC) and multiple glycolytic genes, including glucose transporter SLC2A1, hexokinases (HK2 and HK3), 6-phosphofructo-2-kinase/fructose-2,6-bisphosphatase 3 (PFKFB3), glyceraldehyde-3-phosphate dehydrogenase (GAPDH), phosphoglycerate kinase 1 (PGK1) and beta-enolase (ENO3) (**Figs 3**, **S2E**). Macrophages also increased the expression of anti-apoptotic genes, such as BCL2, BCL2A1, BCL2L11 and BCL2L2, in response to *A. baumannii* infection (**Fig 3, Table S4**). Simultaneously, aerobic metabolism was suppressed, as evidenced by the upregulation of pyruvate dehydrogenase kinase isozyme 1 (PDK1) and downregulation of key oxidative phosphorylation-associated genes, including NADH dehydrogenase [ubiquinone] I (i.e., NDUFA1, NDUFA3, NDUFA5, NDUFA6, NDUFB10, NDUFB4, NDUFC2, NDUFS5 and NDUFS6), NADH ubiquinone oxidoreductase (i.e., ND1, ND2, ND3, ND4, ND4L, ND5 and ND6), cytochrome b (CYTB), cytochrome b-c1 complex (i.e., UQCRHL and UQCRB), cytochrome c oxidase (i.e., COX2, COX3, COX6B1, COX7A2L and COX7B) and ATP synthase subunits (i.e., ATP6, ATP8, ATP5PD, ATP5F1E and ATP5MG) (**Figs 3 and S2F, Table S4**). Interestingly, genes involved in the degradation of HIF-1α via the ubiquitin-dependent proteasomal pathway, including prolyl hydroxylase (EGLN3), E3 ubiquitin ligase complex (RBX1) and elongin-B (ELOB) were also upregulated (log_2_FC values of 5.44, 1.45 and 1.52, respectively; [**Fig 3**]). These findings highlight the activation of HIF-1 signaling and significant metabolic shifts in macrophages upon infection with *A. baumannii* AB5075.

### Impairment of *rcnB*, *ompW*, and *traR/dksA* families diminishes *in vitro* polymyxin B tolerance and reduces bacterial survivability *in vivo*

To elucidate the roles of the key DEGs on polymyxin tolerance in *A. baumannii* when interacting with macrophages, we firstly screened the top DEGs by conducting time-kill kinetics using the AB5075 transposon mutants with single-gene disruption in several key candidate targets. Remarkably, mutants with disruptions in ABUW_RS02965 (*rcnB* family), ABUW_RS02730 (*traR/dksA* C4-type zinc finger), and ABUW_RS06025 (*ompW* family) exhibited substantially enhanced killing by polymyxin B (4 mg/L) than the wild-type strain, with no viable colonies detected at 1 and 5 h (**Fig 4A**). Additionally, disruption of *rpmE2* (type B 50S ribosomal protein L31), *tatA* (Sec-independent protein translocase subunit), and ABUW_RS08590 (universal stress protein) resulted in >5 log_10_ CFU/mL increased killing compared to that of polymyxin-treated wild-type at 1 and 5 h (**Figs 4A, 4B**).

**Figure 4:**
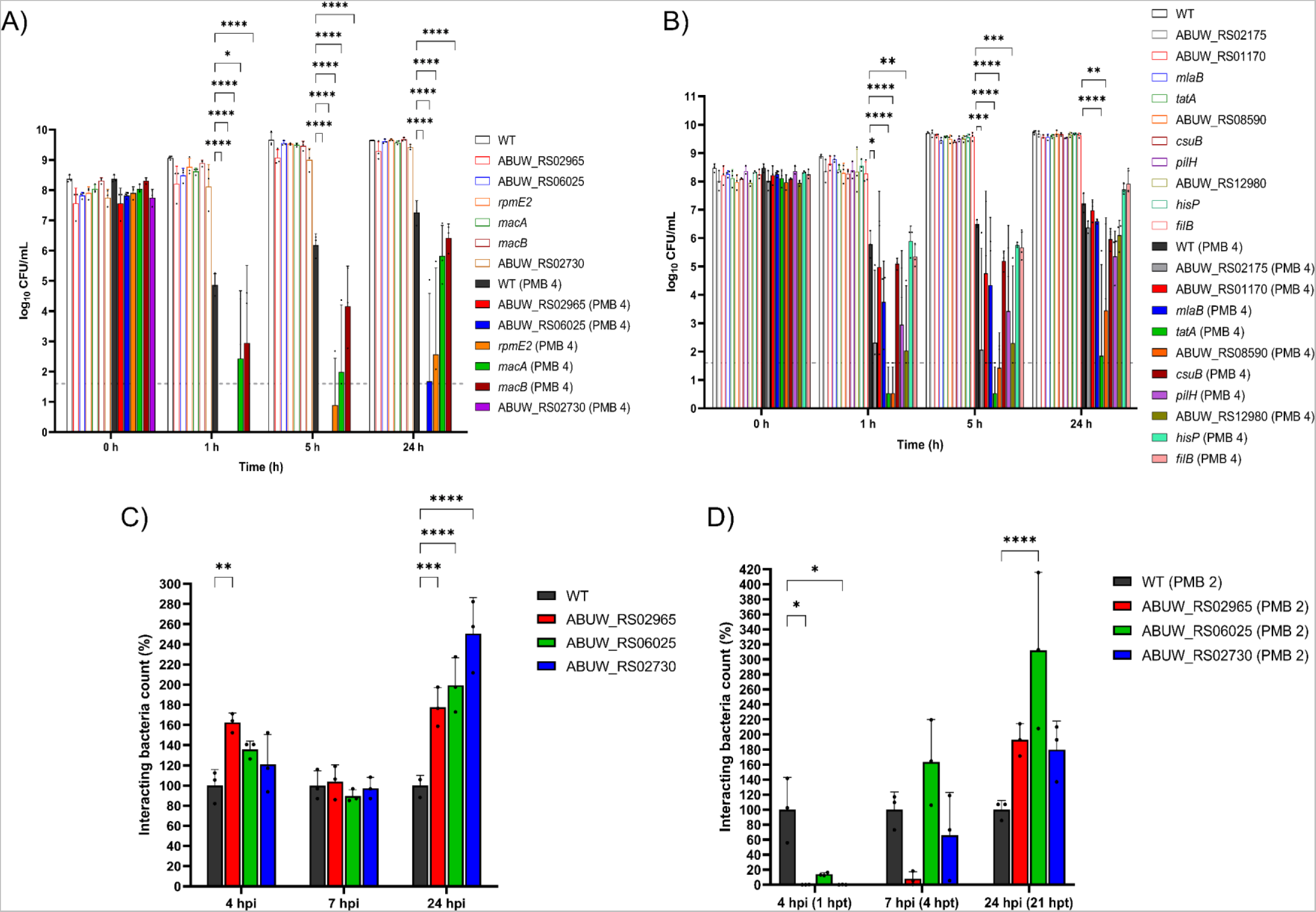
Killing kinetics of polymyxin B against *A. baumannii* AB5075 transposon mutants compared to that of the wild-type (WT) in the absence and presence of macrophages. (**A**) and (**B**) show the time-kill profiles of AB5075 mutants and the WT following the treatment of 4 mg/L polymyxin B (PMB) in the absence of macrophages. Interacting bacterial count of AB5075 mutants and the WT following MOI 1,000 infection on macrophages in the absence (**C**) and presence (**D**) of 2 mg/L polymyxin B treatment. Data are presented as means ± SD of biological triplicate. Two-way ANOVA with Tukey’s HSD post-hoc test, * p <0.05, ** p <0.01, *** p <0.001, **** p <0.0001. Dashed lines represent the limit of detection.

Subsequently, we examined the survivability of the top three bacterial mutants (*rcnB*, *traR/dksA*, and *ompW* family mutants) that demonstrated the greatest reduction in polymyxin tolerance during interaction with macrophages in the presence and absence of polymyxin B. In the absence of polymyxin B treatment, *rcnB* mutant showed 62.5 ± 9.47% (mean ± SD) increase in the interacting bacterial count at 4 h post-infection compared to the wild-type. At 24 h, all three *rcnB*, *traR/dksA*, *ompW* mutants exhibited significantly higher numbers of interacting bacteria with macrophages (around 70% to 150% increase compared to the wild-type) (**Fig 4C**). Notably, in the presence of 2 mg/L polymyxin B, interacting bacteria fractions of *rcnB* and *traR/dksA* mutants were significantly reduced (>99% reduction compared to the polymyxin-treated wild-type, FDR <0.05) at 4 h post-infection (i.e., 1 h post polymyxin B treatment; [**Fig 4D**]). Additionally, although the reduction did not reach statistical significance, the *ompW* mutant showed a notable ~85% reduction (FDR, 0.09) at 1 h post polymyxin B treatment and the *rcnB* mutant demonstrated >90% reduction (FDR, 0.06) at 4 h post polymyxin B treatment (**Fig 4D**).

Furthermore, *in vivo* survivability of *rcnB*, *ompW* and *traR/dksA* mutants were compared to AB5075 wild-type using an immunocompetent mouse bloodstream infection model. The *rcnB* mutant exhibited the lowest bacterial viability at 2 h post-infection, followed by the *traR/dksA* mutant and *ompW* mutant (2.47, 2.24, and 0.69 log_10_ CFU/mL lower than the wild-type, respectively [**Fig 5**]). At 6 h post-infection, viable counts of *rcnB* mutant remained significantly reduced (1.74 log_10_ CFU/mL lower than the wild-type, respectively), while the *ompW* and *traR/dksA* mutants showed no statistically significant difference (**Fig 5**). In the presence of polymyxin treatment, the *in vivo* viability of the *rcnB* mutant significantly decreased, showing a mean reduction of 1.49 log_10_ CFU/mL compared to polymyxin-treated wild-type at 6 h post-infection (i.e., 4 h post-treatment; [**Fig 5**]). Although the adjusted p-value is 0.06, a notable decrease in viable count of polymyxin-treated *traR/dksA* mutant was observed, showing 1.11 log_10_ CFU/mL lower than polymyxin-treated wild-type (**Fig 5**).

**Figure 5:**
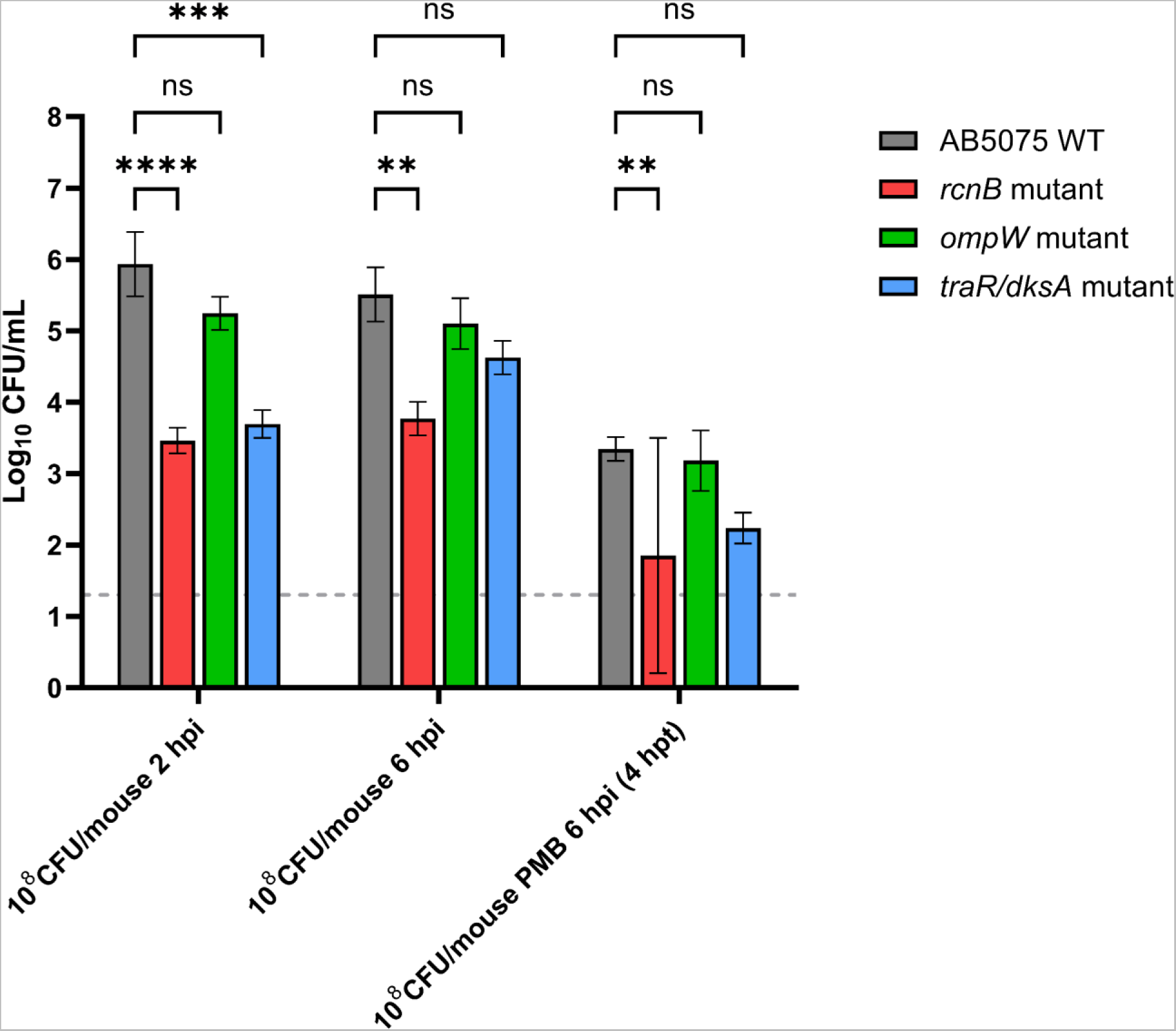
*rcnB* and *traR/dksA* mutants exhibit reduced survivability *in vivo*. Viable bacterial counts of AB5075 WT and transposon mutants with single-gene disruption in *rcnB*, *ompW* and *traR/dksA* families following bloodstream infection in immunocompetent mice in the absence and presence of 4 mg/kg intravenous administration of polymyxin B (PMB). Three biological replicates per experimental condition were utilized. Two-way ANOVA with Dunnett’s multiple comparisons test, * p <0.05, ** p <0.01, **** p <0.0001. Dashed line represents the detection limit (1.30 log_10_ CFU/mL). Hpi, hour post infection; hpt, hour post polymyxin B treatment.

## DISCUSSION

Using our *in vitro* host-pathogen-drug tripartite model [19], we employed dual RNA-sequencing to simultaneously uncover adaptive transcriptional responses in human macrophages and *A. baumannii* during co-culture infection with and without polymyxin B treatment. Our results revealed the significant role of *rcnB*, *ompW*, and *traR/dksA* families in *A. baumannii* and hypoxia-induced signaling in infected macrophages during the interplay among bacterial stress tolerance strategies, macrophages, and polymyxin treatment. Notably, our findings provide new insights on the synergistic disruption on *A. baumannii* stress tolerance by human macrophage and polymyxin B at the transcriptomic level (**Fig 6**). Furthermore, we identified key players in stress sensing and response regulation in *A. baumannii*, such as *hfq*, *baeSR*, *bfmSR*, and *envZ*, along with effector machineries including RND and *macAB* MDR efflux pumps, *betAB* and *otsAB* osmoprotective systems, and oxidative stress-associated genes. Together, these findings offer potential avenues for further therapeutic exploration and illuminate the mechanisms of polymyxin antibacterial activity and *A. baumannii* pathogenesis in human macrophages.

**Figure 6:**
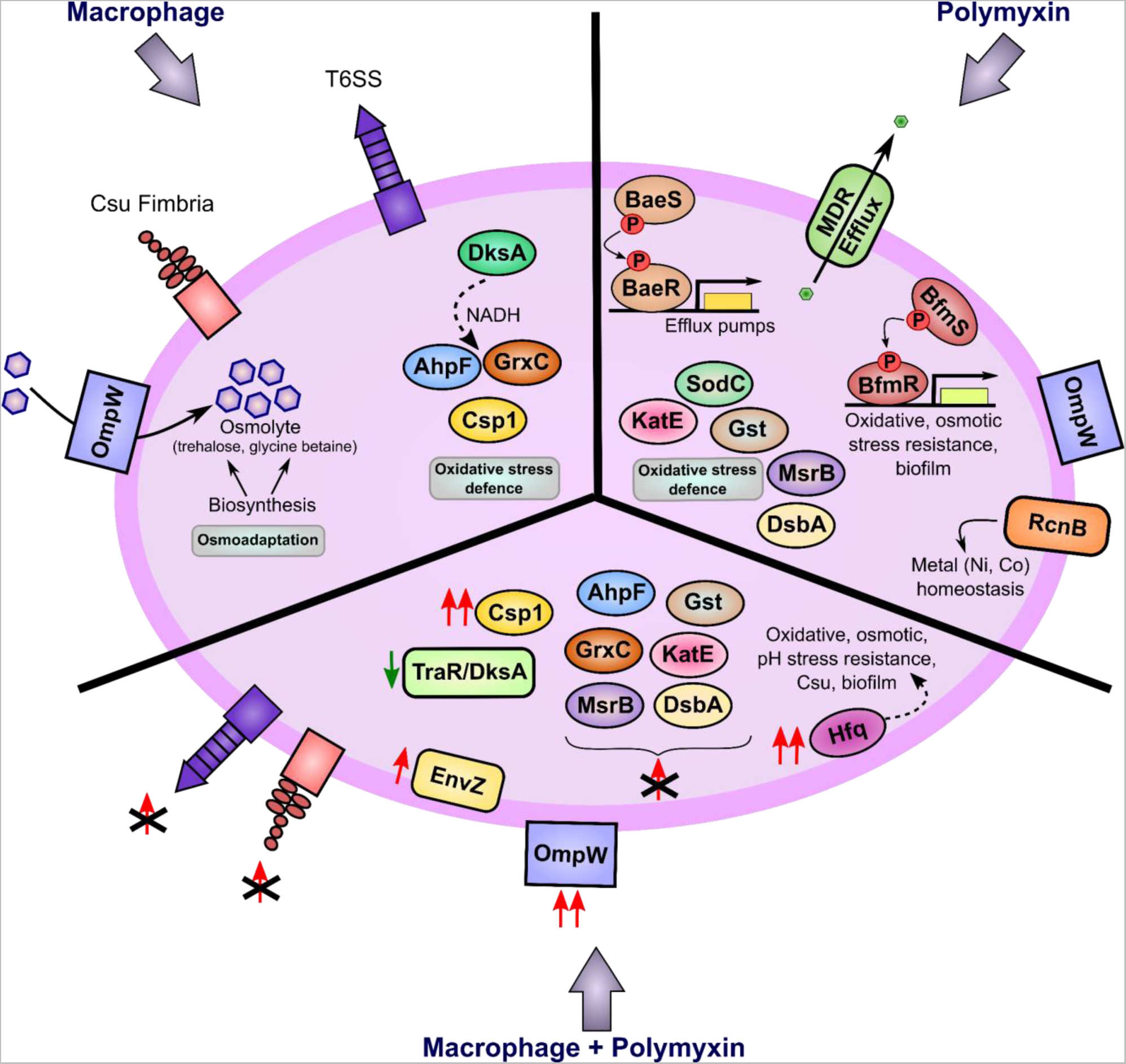
Schematic summary of differential transcriptional regulation and perturbation of key stress tolerance machineries in *A. baumannii* AB5075 in response to macrophages, polymyxin B, and the combination. Ni, nickel; Co, cobalt.

In infected macrophages, polymyxin treatment caused overexpression of the envelope stress response associated RNA chaperone *hfq* in *A. baumannii* (**Fig 1A**). The riboregulator *hfq* plays a crucial role in facilitating the efficient base pairing of small regulatory RNAs (sRNAs), which stabilizes and/or activates mRNA translation [20]. These actions establish a complex post-transcriptional network that regulates the diverse environmental adaptation and virulence strategies of bacteria [20]. It has been reported that the loss of *hfq* function in *A. baumannii* ATCC 17978 is associated with reduced stress tolerance (i.e., oxidative, osmotic, acidic and alkaline pH stresses), impaired *csu* fimbrial biogenesis, decreased biofilm formation, compromised adhesion and invasion of host epithelial cells, and diminished intramacrophage survival [21]. Furthermore, under the tripartite condition in the present study, *tatAC* from the twin-arginine translocation (Tat) system and *lptG* from the lipopolysaccharide (LPS) transport system were significantly upregulated. The bacterial Tat system is well known in facilitating the transport of membrane proteins and export of virulence factors (e.g. phospholipases, elastase, alkaline phosphatase, and metal-acquisition proteins) [22, 23]; therefore, our findings indicate that bacterial attempts to facilitate membrane repair and enhance pathogenicity in response to polymyxin treatment in macrophages (**Fig 2**).

In *A. baumannii* the upregulation of bacterial envelope stress sensing and signal transduction systems, *baeSR* and *bfmSR,* reveals their vital roles in the bacterial response to polymyxin-induced outer membrane damage (**Fig 1D**) [24]. The *baeSR* system regulates the expression of multidrug efflux pumps, including *adeABC, adeIJK* and *macAB-tolC* [25–28]. Notably, our study observed overexpression of *macAB*, ABUW_RS16345 (*adeC/adeK/oprM* family multidrug efflux complex outer membrane factor) and ABUW_RS04525 (RND transporter permease) across all the polymyxin-treated *A. baumannii* samples, and the disruption in *macA* resulted in enhanced polymyxin susceptibility (**Figs 1A and 4A**). Moreover, while the presence of macrophages in the tripartite condition abrogated the polymyxin-induced *baeR* overexpression, the associated RND and *macAB* efflux systems remained overexpressed, suggesting compensatory cross-regulation within the *baeR* regulon by other two-component systems. Taken together, these findings underscore the critical role of *baeSR* and multidrug efflux systems, such as *macAB-tolC*, in mediating polymyxin B tolerance.

In response to polymyxin B treatment and/or macrophage infection, *A. baumannii* also underwent significant osmoadaptation. Notably, the *ompW* family gene ABUW_RS06025 was consistently overexpressed across all treatment groups, with an additive overexpression observed in the tripartite condition (AB+THP+PMB) (**Fig 1C**). In line with this, the interacting bacteria in the tripartite condition exclusively overexpressed *envZ*, which encodes the membrane-bound sensor kinase of the *envZ/ompR* two-component system involved in osmotic stress response by regulating OMP porin expression (**Figs 1C, 1D**) [29]. These observations indicate bacterial attempts to counteract greater envelope and osmotic stresses in the tripartite condition. Supporting our speculation, OmpW porin protein of *Vibrio cholerae* has been found to transport carnitine and other compatible solutes, playing a crucial role in osmoadaptation during growth under hypersaline conditions [30]. Moreover, the crucial role of the *ompW* family in polymyxin tolerance is further supported by the reduced tolerance observed in *A. baumannii* transposon mutant with defective ABUW_RS06025 (**Fig 4**). Additionally, the upregulation of trehalose biosynthetic enzymes *otsA* (ABUW_RS15165; trehalose-6-phosphate [T6P] synthase) and *otsB* (trehalose phosphatase) reflects the strategy of *A. baumannii* to cope with osmotic stress caused by polymyxin-induced envelope damage. In response to macrophages, bacterial choline catabolic genes *betAB* were transcriptionally upregulated to mediate glycinebetaine biosynthesis as an osmoprotectant. The accumulation of glycinebetaine suppresses the intake and biosynthesis of other intracellular osmolytes, including trehalose, which explains the downregulation of T6P synthase *otsA* specifically in the infection condition (AB+THP) [31–33]. The cooperative role of macrophages and polymyxin B in perturbing bacterial osmo-metabolism is highlighted by the abrogation of overexpression in osmoprotectant trehalose and glycinebetaine biosynthesis in the tripartite condition (**Fig 1A**). Significantly, the deletion of *betAB* in *P. aeruginosa* has demonstrated a decrease in bacterial survival in a murine pneumonia model, highlighting the essential role of choline catabolism in bacterial persistence within the host [34]. Bacterial osmoadaptation in mediating tolerance towards polymyxin and macrophages is underscored by these collective findings.

Our study reaffirms the critical roles of antioxidant mechanisms and stringent stress regulatory systems in *A. baumannii* in coping with the stress induced by polymyxin and macrophage infection. Polymyxin B exposure triggers the activation of ROS-scavenging enzymes, including catalases (*katE*, ABUW_RS12185) and superoxide dismutase (*sodC*), aligning well with our previous studies [19, 35, 36]. Polymyxin-induced membrane damage disrupts crucial bacterial protein folding systems, triggering the upregulation of the *dsb* system for proper disulfide bond formation, protein stabilization, and virulence factor assembly, along with the methionine sulfoxide reductase (*msr*) system that repairs oxidized methionine residues [37, 38]. Glutathione-based antioxidant systems are employed by *A. baumannii* to counteract polymyxin-induced oxidative stress, as evidenced by the overexpression of glutathione peroxidases (ABUW_RS11455 and *btuE* [ABUW_RS18150]) to breakdown hydrogen peroxide, glutathione *S*-transferases (*gst*, *yfcF*, ABUW_RS18250) to bind and detoxify exo- and endogenous electrophiles, and glutathione-dependent disulfide-bond oxidoreductase *yghU* to target organic hydroperoxides [39, 40]. Moreover, after infecting macrophages *A. baumannii* cells exhibited increased expression of stress adaptation proteins involved in transport of sulfate and taurine as antioxidative precursors, *csp1* for oxidative and acidic pH stress adaptation, glutaredoxin for thiol-disulfide oxidoreduction, and *ahpF* for peroxide scavenging [19, 41–44].

Notably, infecting *A. baumannii* exhibits increased expression of *dksA* (**Fig 1A**), a pivotal regulator of central carbon metabolism responsible for generating essential reducing equivalents like NAD(P)H, which fuel thiol-based antioxidant systems such as the glutathione-based redox system and alkyl hydroperoxide reductase [45]. Our previous findings showed that transposon disruption of *dksA* reduced *A. baumannii* survival within macrophages, highlighting the vital role of *dksA*-mediated antioxidant activity in resisting macrophage oxidative burst [19]. TraR, a distant protein homolog of DksA, exhibits global regulatory effects of both DksA and stress alarmone (p)ppGpp in the presence of various nutritional stresses for survival [46]. Particularly, we observed a specific downregulation of ABUW_RS02730 (*traR/dksA* C4-type zinc finger) in the tripartite condition, suggesting a potential collaboration between polymyxin and macrophages in perturbing such stringent stress response machinery in *A. baumannii* (**Fig 1A**). The therapeutic potential of targeting this stringent response system was further supported by our transposon mutant study, where disruption of this gene significantly enhanced polymyxin antibacterial efficacy (**Figs 4A, 4D**).

Another important finding of our study is that *A. baumannii* showed a striking upregulation of multiple genes from the *rcnB* family, involved in nickel/cobalt homeostasis, upon polymyxin treatment irrespective of the presence or absence of macrophages (**Fig 1B**). Disruption of ABUW_RS02965, a member of the *rcnB* family, significantly diminished bacterial tolerance to polymyxin B *in vitro* and *in vivo*, and reduced virulence *in vivo* (**Figs 4A, 4D, 5**). This underscores, for the first time, the pivotal role of *rcnB* in polymyxin tolerance and bacterial virulence. While the exact function and mechanism of the *rcnB* family remain unclear, previous research suggests that *rcnB* likely modulates *rcnA*-mediated metal efflux to maintain intracellular nickel and cobalt ion pools [47]. Nickel and cobalt are essential for the catalytic activity of various enzymes involved in hydrogen, nitrogen, and carbon metabolism [48, 49]. For instance, nickel acts as a catalytic cofactor in enzymes, including [NiFe]-hydrogenase, which reversibly converts molecular hydrogen to electrons and protons; urease, which is involved in nitrogen recycling; and carbon monoxide dehydrogenase, which is involved in carbon recycling [48]. Cobalt is also involved in diverse biological processes, either in the form of vitamin B12 or as a component of non-corrin cobalt proteins such as glucose isomerase which converts glucose to fructose, and methionine aminopeptidase which regulates protein turnover [49]. Impairment of the *rcnB* family may disrupt the control of these metal ion pools and downstream enzymatic regulation, leading to compromised stress tolerance of *A. baumannii* against polymyxins and host immune defenses.

*A. baumannii* strategically regulated its iron uptake and assimilation systems in response to polymyxin B and macrophage infection, reducing iron availability to minimize ROS production via the Fenton reaction (**Fig 1B**). Meanwhile, infected macrophages upregulated metal transporter SLC39A14 and transferrin receptors to enhance uptake of manganese, zinc, cadmium, and other metals, contributing to nutritional immunity against invading bacteria (**Fig 3**) [50]. Interestingly, the downregulation of host serotransferrin (TF) suggests macrophages sequestered available metal-bound complexes instead of synthesizing their own transferrin, consistent with our previous proteomic findings of increased lactotransferrin (LTF) and heme scavengers (i.e., albumin, apolipoprotein, haptoglobin) in infected macrophages [19]. Furthermore, the observed downregulation of heme catabolic HMOX1 in infected macrophages is consistent with our previous proteomics study, suggesting the host attempt to elevate pro-oxidant and pro-inflammatory free heme levels for immune activation [19]. Heme is known to induce TLR-triggered TNF-alpha production, NF-ĸB and mitogen-activated protein kinase (MAPK)-mediated cytokine production, G protein-coupled receptor (GPCR)-mediated neutrophil recruitment, oxidative burst in phagocytes, and delayed neutrophil apoptosis [51–54]. The observed upregulation of TLR2, NOXA1, and the enrichment of TNF, NF-κB, MAPK, and phosphatidylinositol-3-kinase (PI3K)-AKT signaling cascades in infected macrophages were likely triggered by elevated levels of free heme (**Fig S2E**). These pro-inflammatory responses, such as TNF and IL-6 signaling, would subsequently activate the coagulation cascade, leading to bacterial entrapment within fibrin clots (**Fig 3**) [55].

Importantly, our findings highlight the pivotal role of HIF-1 signaling in *A. baumannii*-infected macrophages, promoting host survival and adaptation to low oxygen conditions through metabolic regulation. During microbial infection, various factors (e.g. infection-induced hypoxia) and pathological stressors (e.g. bacterial LPS and siderophores) can trigger the stabilization and activation of HIF-1α. This, in turn, leads to the activation of immune and inflammatory responses against the infection [56, 57]. The upregulation of vascular endothelial growth factors in infected macrophages is a hallmark of HIF-1α activation, facilitating oxygenation and glycolysis to anaerobically generate energy to sustain macrophage survival (**Fig 3**) [58–60]. The observed upregulation of HIF-1α degradation via ubiquitin-mediated pathways may be a bacterial strategy to counteract the heightened HIF-1 signaling in macrophages (**Fig 3**). The transcriptional downregulation of bacterial oxidative phosphorylation during macrophage infection, including NADH-quinone oxidoreductases and cytochrome bd oxidases, likely indicates the bacterial attempt to increase the availability of molecular oxygen (**Table S1**). This, in turn, triggers the activity of host prolyl hydroxylase EGLN3 (Fe^2+^- and oxygen-dependent) to hydroxylate specific proline residues of HIF-α and subsequently target it for degradation [55]. Interestingly, infected macrophages simultaneously downregulated their aerobic metabolism, as indicated by the upregulation of pyruvate dehydrogenase kinase isozyme 1 (PDK1) that hinders pyruvate conversion into acetyl-CoA, thereby reducing metabolic flux into the TCA cycle [61]. Furthermore, the downregulation of multiple genes involved in oxidative phosphorylation, including those in the electron transport chain and ATP synthase subunits, in infected macrophages further decreased aerobic energy production (**Fig 3**). This metabolic shift towards glycolysis aligns well with the pro-inflammatory (M1) macrophage phenotype, while anti-inflammatory (M2) macrophages rely more on oxidative phosphorylation [62].

## CONCLUSIONS

The interplay between host and pathogen stress adaptations shapes infection outcomes and treatment efficacy. Our study investigated the intricate regulatory networks and downstream effectors that drive the unique stress adaptation responses of *A. baumannii* and macrophages during infection and polymyxin treatment. The transcriptomic signatures specific to infection, polymyxin B treatment, and the tripartite host-pathogen-drug conditions primarily revolve around membrane homeostasis and stress response machineries, encompassing redox, osmotic, and nutritional (metal) stress responses. Importantly, we demonstrate for the first time the synergistic impact of macrophages and polymyxin B on perturbing stress responses in *A. baumannii* at the transcriptomic level; and reveal potential targets for the discovery of new antibiotics.

## MATERIALS AND METHODS

### Bacterial strains, media and antibiotic

MDR *Acinetobacter baumannii* AB5075 wild-type (WT) is originally a wound isolate from a patient at the Walter Reed Army Medical Center [63]. The tetracycline-resistant AB5075-UW T26 transposon mutants with single-gene disruption used in this study (**Table S5**) were obtained from the University of Washington [64]. To prepare logarithmic-phase bacterial cultures, −80°C frozen stocks of AB5075 WT and the mutants were streaked onto nutrient agar (NA) and Luria-Bertani (LB) agar, with the latter supplemented with 10 mg/L tetracycline hydrochloride (Cat# T7660-5G; Sigma). Following aerobic incubation at 37°C for 18 h, a single bacterial colony was inoculated into cation-adjusted Mueller-Hinton broth (CaMHB) and grown at 37°C with shaking (200 rpm) for 18 h. A 100-fold dilution of the overnight cultures was performed in pre-warmed CaMHB followed by further incubation at 37°C (200 rpm) to reach logarithmic-phase (approximate optical density [OD] of 0.5 at 600 nm). Sterile stocks of polymyxin B sulphate (PMB; Cat# 86-40302, Betapharma, Shanghai, China) were prepared in Milli-Q water (Millipore, USA) with filter sterilization using a 0.22-µm syringe filter (Sartorius, Germany).

### Mammalian cell culture

Human monocytic cell line THP-1 (ATCC TIB-202) was cultured in eukaryotic growth media Roswell Park Memorial Institute (RPMI) 1640 medium with additional supplementation of 25 mM 4-(2-hydroxyethyl) piperazine-1-ethanesulfonic acid (HEPES) buffer and 10% fetal bovine serum (FBS; Lot# 15703, Bovogen, Australia) and grown at 37°C (5% CO_2_). Differentiation of THP-1 cells into macrophage-like cells were conducted via treatment with 25 nM of phorbol 12-myristate 13-acetate (PMA; Santa Cruz Biotechnology, USA). After 48-h PMA stimulation, treatment media was replaced with fresh pre-warmed RPMI 1640 media supplemented with 25 mM HEPES and 10% FBS, followed by at least 24-h cell resting in PMA-free media before conducting any experiment.

### Dual RNA-seq sample preparation

The *in vitro* host-pathogen-drug co-culture conditions employed in this study were previously described in detail in our earlier publication [19]. Briefly, macrophages were infected by *A. baumannii* AB5075 at a multiplicity of infection (MOI) of 1,000 for 4 h in RPMI 1640 media supplemented with 25 mM HEPES and 10% heat-inactivated FBS. For antibiotic-treated groups, 30 mg/L of polymyxin B was added at 3 h post infection for another 1 h. All experiments (including bacterial samples without macrophages) were performed in RPMI 1640 media supplemented with 25 mM HEPES and 10% heat-inactivated FBS in 6-well plates (Corning; Jiangsu Province, China) at 37°C and 5% CO_2_. At 4 h, the cell monolayer was washed twice with ice cold 1×Dulbecco’s phosphate-buffered saline (DPBS; Gibco, Paisley, UK) for removal of non-interacting extracellular bacteria. Bacterial samples without macrophages were centrifuged at 10,000 × *g* for 5 min at 4°C to pellet bacteria then washed twice with ice cold 1×DPBS. All cell samples were digested using TRIzol reagent (CAT# 15596018, Thermo Fisher Scientific) before proceeding to RNA isolation. Three biological replicates were performed across all experiments.

### RNA isolation, dual-RNA-seq and bioinformatic analysis

Total RNA isolation was performed using a Direct-zol RNA MiniPrep Plus kit (Zymo Research; CAT# R2072). RNA samples were sent to Genewiz Genomic Center (China) for sample quality assessment, clean-up, cDNA library preparation and sequencing using Illumina HiSeq. Briefly, the sequencing library was prepared using NEBNext Ultra RNA Library Prep Kit for Illumina. Poly(A) mRNA was isolated via NEBNext Poly(A) mRNA Magnetic Isolation Module (NEB) or the Ribo-Zero rRNA removal Kit (Illumina). The mRNA was fragmented and primed using NEBNext First Strand Synthesis Reaction Buffer and NEBNext Random Primers. Synthesis of first and second strand cDNAs were carried out using ProtoScript II Reverse Transcriptase and Second Strand Synthesis Enzyme Mix, respectively, followed by purification of the double-stranded cDNA using AxyPrep Mag PCR Clean-up (Axygen). Double-stranded cDNA that underwent end repair and dA-tailing by End Prep Enzyme Mix was then proceeded to adaptor ligation to both ends and size-selected using AxyPrep Mag PCR Clean-up (Axygen) to recover fragments of approximately 360 base pairs (bp). Following polymerase chain reaction (PCR) amplification using P5 and P7 primers, the PCR products were purified by AxyPrep Mag PCR Clean-up (Axygen). Quality assessment and quantification of the PCR products were respectively performed using the Agilent 2100 Bioanalyzer system (Agilent Technologies, Palo Alto, CA, USA) and a Qubit 2.0 Fluorometer (Invitrogen, Carlsbad, CA, USA). The multiplexed libraries were sequenced by Illumina HiSeq instrument (Illumina, San Diego, CA, USA) on a 2×150 bp paired-end run. Image analysis and base calling were carried out using the HiSeq Control Software (HCS)+OLB+GAPipeline-1.6 (Illumina).

The raw sequence reads of AB5075 and THP-1 macrophage were aligned with the respective reference genome sequence downloaded from the NCBI Assembly database using RSubread [65]. The RNA sequence data from biological triplicates were analyzed using the voom and limma linear modeling methods via Degust interactive Web-based RNA-seq visualization software (http://www.vicbioinformatics.com/degust/) [66]. Statistically significant DEGs were defined as those with log_2_FC >1 or <-1 and FDR <0.05. The functions of bacterial DEGs were analyzed based on Clusters of Orthologous Groups (COGs) and Kyoto Encyclopedia of Genes and Genomes (KEGG) pathways using R, with the pathway enrichment statistical cut-off of FDR <0.2 using Fisher’s Exact Test. Host DEGs were analyzed on WebGestalt to identify the enriched KEGG pathways (Benjamini-Hochberg; FDR <0.05) [67].

### *In vitro* investigation of polymyxin antibacterial activity against AB5075 mutants in the absence and presence of THP-1 macrophages

To examine the time-kill kinetics of polymyxin B against AB5075 wild-type and the transposon mutants, bacteria from respective logarithmic-phase cultures in CaMHB were firstly pelleted by centrifugation at 3,220 × *g* for 20 min. Bacterial pellet was resuspended in pre-warmed RPMI 1640 media supplemented with 25 mM HEPES and 10% heat-inactivated FBS to prepare a starting culture of ~10^8^ CFU/mL. Polymyxin B was added to respective treatment groups to achieve 4 mg/L, followed by incubation at 37°C (200 rpm). At baseline (0 h) and 1, 5 and 24 h post antibiotic treatment, bacterial viability was assessed using spot assay (25 μL per spot; the limit of detection was 1.60 log_10_ CFU/mL) of appropriately diluted bacterial cultures onto nutrient agar followed by incubation at 37°C for 16–18 h prior to colony counting.

For the *in vitro* infection studies, macrophages were infected at MOI 1,000 on 24-well plates and incubated at 37°C and 5% CO_2_. At 3 h post infection, non-interacting extracellular bacteria were aspirated, and the macrophage cell monolayer washed twice with pre-warmed 1×DPBS. Pre-warmed RPMI 1640 media supplemented with 25 mM HEPES and 10% heat-inactivated FBS containing 2 mg/L polymyxin B was added to the respective treatment wells prior to further incubation with 5% CO_2_ at 37°C. At 1, 4 and 21 h post polymyxin B treatment, cell monolayer was washed twice with ice cold 1×DPBS to remove any non-interacting extracellular bacteria. Lysis of macrophages was conducted using 1% Triton X-100 for 1 h on ice to release the interacting bacteria from infected macrophages into the lysate. Bacteria in the macrophage lysates were pelleted by centrifugation at 10,000 × *g* (4°C) for 10 min and subsequently resuspended in 0.9% NaCl solution. Following serial dilutions, 25 μL of bacterial suspension was spot-plated onto nutrient agar and incubated at 37°C for 16 - 18 h before colony counting. The viable bacterial counts were statistically analyzed using two-way ANOVA and Tukey’s HSD post-hoc test (GraphPad Prism 9.2.0).

### Mouse infection model studies

All animal experiments were approved by the Monash Animal Ethics Committee and conducted in concordance with the Australian Code of Practice for the Care and Use of Animals for Scientific Purposes. A non-neutropenic mouse bloodstream infection model using female Swiss mice (8-week-old, body weight between 22.3 to 34.0 g) was employed to compare virulence of AB5075 transposon mutants with wild-type strain (Monash University Clayton Campus, Victoria, Australia). Mice were housed with water and food provided *ad libitum* (Monash University Clayton Campus). To establish bloodstream infection of 10^8^ CFU/mouse, 100 µL of 10^9^ CFU/mL early logarithmic-phase bacterial suspension of AB5075 wild-type (WT) or its transposon mutants with single-gene disruptions in ABUW_RS02965 (*rcnB* family), ABUW_RS06025 (*ompW* family) and ABUW_RS02730 (*traR/dksA* C4-type zinc finger) was administered to each mouse via tail vein. At 2 h post-inoculation, the treatment group received intravenous administration of polymyxin B at a dosage of 4 mg/kg. Following 2 h and 6 h infection, mice were euthanized with inhaled carbon dioxide prior to blood collection via cardiac puncture. Blood samples were serially diluted in sterile 0.9% NaCl solution followed by spiral plating (50 µL) on nutrient agar for overnight incubation (37°C, 16– 18 h) prior to colony counting (limit of detection, 1.30 log_10_ CFU/mL). Three mice were used in each experimental condition and the mice were assigned through randomization. Viable bacterial counts were statistically analyzed using two-way ANOVA with Dunnett’s multiple comparisons test (GraphPad Prism 9.2.0).

## ACKNOWLEDGEMENTS AND FUNDING

We thank Dr. Jinxin Zhao for his assistance in liaising with GENEWIZ for the sample processing and sequencing. We thank Dr. Jiping Wang and Mr. Ke Chen for conducting mice experiments. This research was funded by the National Institute of Allergy and Infectious Diseases of the National Institutes of Health (R01 AI146160). The content is solely the responsibility of the authors and does not necessarily represent the official views of the National Institute of Allergy and Infectious Diseases or the National Institutes of Health. Z.Y.K. was supported by Monash Graduate Scholarship. J.L. is an Australian National Health and Medical Research Council (NHMRC) Principal Research Fellow (APP1157909) and T.N. is an Australian Research Council Future Fellow (FT170100313).

## AUTHOR CONTRIBUTIONS

Conceptualization, J.L.; Experimental design, Z.Y.K., M.A.K.A., J.L.; Technical advice, J.L., M.A.K.A., M.-L.H., Y.Z., T.V., T.Q.Z., and T.N.; Investigation, Z.Y.K.; Data analysis, Z.Y.K., Y.Z.; Writing, Z.Y.K.; Manuscript revision, all authors; Resources, J.L. and T.N.; Funding acquisition, J.L. and Q.T.Z.

## COMPETING INTEREST

All authors declare no competing interests related to this study.

J.L. received grants, speaking honoraria, and consulting fees from Northern Antibiotics, Avexa, Genentech, Healcare, CTTQ, Aosaikang, Jiayou Medicine, MedCom, Fansheng Biotech, DanDi BioScience, Qpex Biopharma, and Xellia Pharmaceuticals. J.L. is CEO of Sliabx Pharmaceuticals and Cinfinno Biotech, and holds equities in both companies.

## LIST OF FIGURE AND TABLE LEGENDS

**Figure S1:**
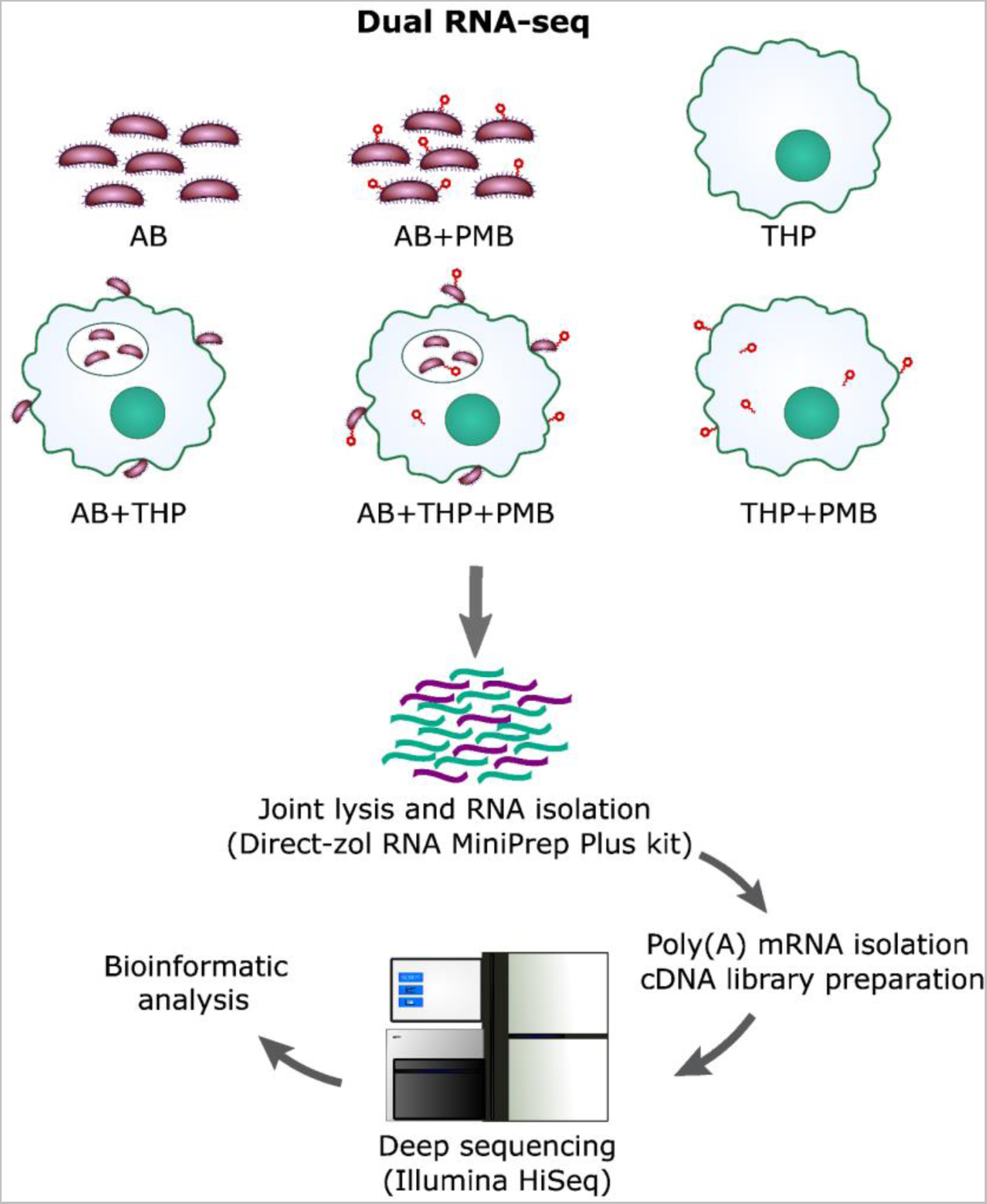
Experimental workflow of transcriptomic analysis of *A. baumannii* AB5075 (AB)-macrophage-polymyxin B (PMB) tripartite axis using a dual RNA-sequencing approach.

**Figure S2:**
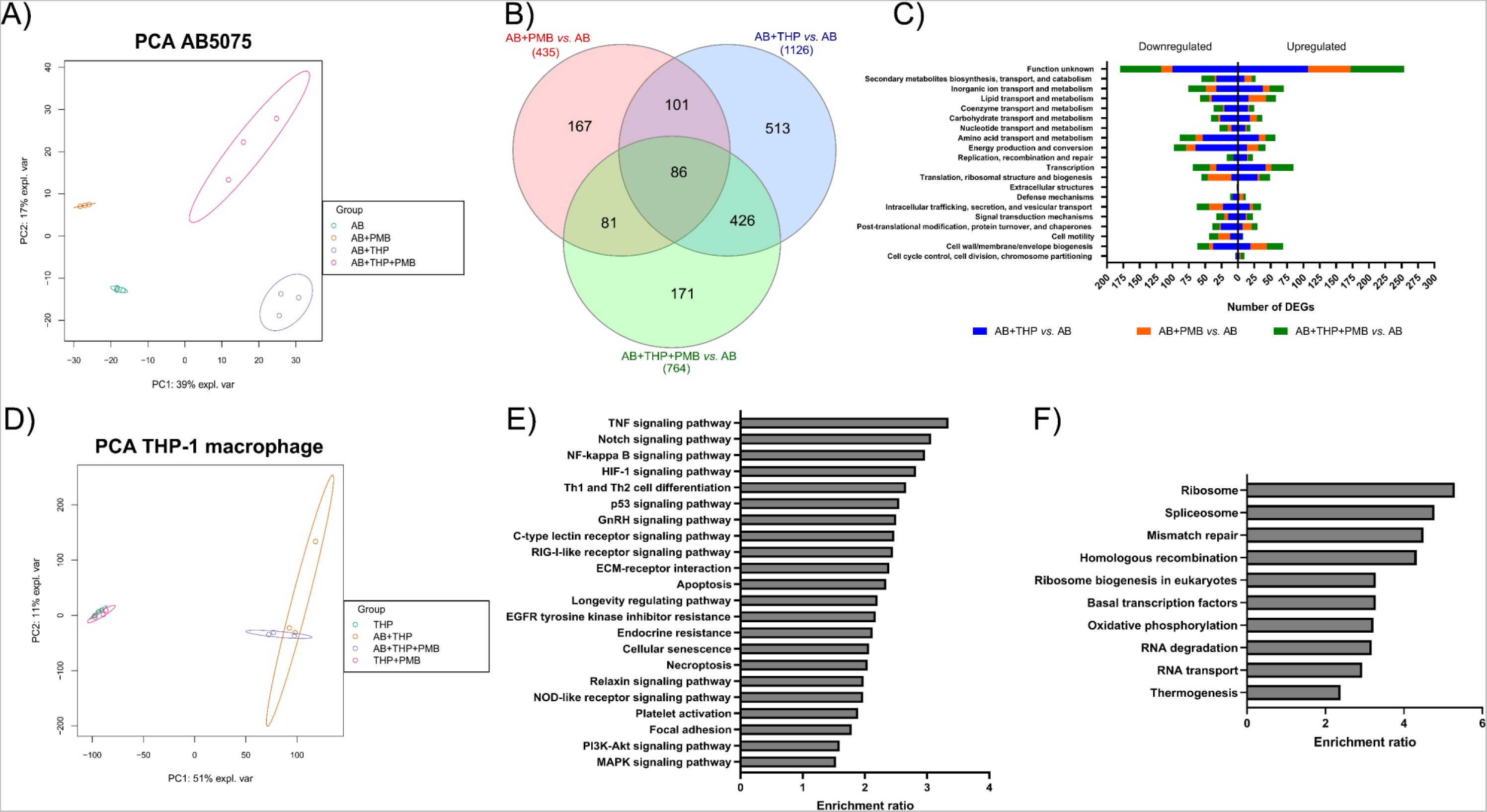
Global transcriptomic changes in *A. baumannii* AB5075 (**A-C**) and macrophages (**D-F**) following different treatments. (**A**) and (**D**) show principal component analysis (PCA) plots of respective expression datasets of AB5075 and macrophages. Venn diagram (**B**) and COG analysis (**C**) of AB5075 DEGs following infection of macrophages (AB+THP), polymyxin B treatment (AB+PMB), and the combination (AB+THP+PMB) compared to untreated bacteria controls (log_2_FC >1 or <-1, FDR <0.05). KEGG pathway enrichment analysis of upregulated (**E**) and downregulated (**F**) macrophage DEGs following AB5075 infection (AB+THP *vs.* THP) using Benjamini-Hochberg FDR <0.05. AB, *A. baumannii* AB5075; THP, THP-1 differentiated macrophages; PMB, polymyxin B; DEGs, differentially expressed genes.

**Table S1.** List of all DEGs in interacting *A. baumannii* AB5075 following infection of macrophages (AB+THP) compared to the untreated bacterial control (log_2_ Fold Change [FC] >1 or <-1, FDR <0.05) and their corresponding gene expression changes in log_2_FC. DEGs, differentially expressed genes; AB, wild-type *A. baumannii* AB5075; PMB, polymyxin B; THP, macrophage-like THP-1 cells.

**Table S2.** List of all DEGs in *A. baumannii* AB5075 following polymyxin B treatment (AB+PMB) compared to the untreated bacterial control (log_2_ Fold Change [FC] >1 or <-1, FDR <0.05) and their corresponding gene expression changes in log_2_FC. DEGs, differentially expressed genes; AB, wild-type *A. baumannii* AB5075; PMB, polymyxin B; THP, macrophage-like THP-1 cells.

**Table S3.** List of all DEGs in interacting *A. baumannii* AB5075 following polymyxin B in the presence of macrophages (AB+THP+PMB), compared to the untreated bacterial control (log_2_ Fold Change [FC] >1 or <-1, FDR <0.05) and their corresponding gene expression changes in log_2_FC. DEGs, differentially expressed genes; AB, wild-type *A. baumannii* AB5075; PMB, polymyxin B; THP, macrophage-like THP-1 cells.

**Table S4.** List of all DEGs in macrophages following infection with *A. baumannii* AB5075 (AB+THP) compared to the untreated host control (log_2_ Fold Change [FC] >1 or <-1, FDR <0.05) and their corresponding gene expression changes in log_2_FC. DEGs, differentially expressed genes; AB, wild-type *A. baumannii* AB5075; PMB, polymyxin B; THP, macrophage-like THP-1 cells.

**Table S5.** List of *A. baumannii* AB5075-UW T26 transposon mutants used in functional studies

